# Intratumoral and extratumoral synapses are required for glioblastoma progression in *Drosophila*

**DOI:** 10.1101/2021.10.14.464400

**Authors:** María Losada-Pérez, Mamen Hernández García-Moreno, Sergio Casas-Tintó

**Author notes:** Corresponding author SC-T.

## Abstract

Glioblastoma (GB) is the most aggressive, lethal and frequent primary brain tumor. It originates from glial cells and is characterized by rapid expansion through infiltration. GB cells interact with the microenvironment and healthy surrounding tissues, mostly neurons and vessels. GB cells project tumor microtubes (TMs) that contact with neurons and exchange signaling molecules related to Wingless/WNT, JNK, Insulin or Neuroligin-3 pathways. This cell to cell communication promotes GB expansion and neurodegeneration. Moreover, healthy neurons form glutamatergic functional synapses with GB cells which facilitate GB expansion and premature death in mouse GB xerograph models. Targeting signaling and synaptic components of GB progression may become a suitable strategy against glioblastoma. In a *Drosophila* GB model, we have determined the post-synaptic nature of GB cells with respect to neurons, and the contribution of post-synaptic genes expressed in GB cells to tumor progression. In addition, we document the presence of intratumoral synapses between GB cells, and the functional contribution of pre-synaptic genes to GB calcium dependent activity and expansion. Finally, we explore the relevance of synaptic genes in GB cells to the lifespan reduction caused by GB advance. Our results indicate that both presynaptic and postsynaptic proteins play a role in GB progression and lethality.

## Introduction

Glioblastoma (GB) is the most lethal and aggressive tumor of the Central Nervous System. GB has an incidence of 3/100,000 adults per year (TAMIMI and JUWEID, 2017), and accounts for 52% of all primary brain tumors. GB originates from glial cells or glial progenitors and causes death within 16 months after diagnosis (Bi and Beroukhim, 2014) due to the low efficacy of standard treatments such as chemotherapy, radiotherapy or surgical resection.

In the last decade, *Drosophila melanogaster* has emerged as a reliable *in vivo* GB model that reproduces the features of human GB (Jarabo et al., 2021; Kegelman et al., 2017; Portela et al., 2019a; Read, 2011; Read et al., 2013). The GB condition is experimentally elicited by the expression of constitutively active forms of EGFR (Epidermal Growth Factor Receptor) and PI3K (Phosphoinositide 3-kinase) in glial cells, which are the two most common mutations in patients (Read et al., 2009). This experimental model has been previously used to study the contribution of RIO kinases (Read et al., 2013), vesicle transport (Portela et al., 2019a), the human kinase STK17A orthologue (Drak) (Chen et al., 2019; Lathia, 2019), and several metabolic pathways in GB (Chi et al., 2019). Consequently, *Drosophila* model of GB is well characterized and suitable to study cellular properties of GB *in vivo*.

Tumor microenvironment and the communication between tumoral cells and neurons are crucial for GB progression and patient survival (Casas-Tintó and Portela, 2019; Jarabo et al., 2021; Orgazy et al., 2014; Portela et al., 2019b, 2020; Qu et al., 2019). In addition, neuronal activity can also stimulate GB growth. Activity-dependent release of neuroliglin-3 (NLGN3) is required for GB progression in xenograft models, and NLGN3 induces the expression of synaptic proteins in glioma cells (Venkatesh et al., 2017). Moreover, GB samples show synaptic gene enrichment (Venkatesh et al., 2019) and glioma cells form functional glutamate synapsis with neighboring neurons, where GB cells are post-synaptic (Venkataramani et al., 2019; Venkatesh et al., 2019). These studies also demonstrated that the pharmacological or genetic inhibition of these electrical signals reduces growth and invasion of the tumor (Venkataramani et al., 2019; Venkatesh et al., 2019).

Synapses are the functional units which underlie animal behavior, memory and cognition. Chemical synapses are specialized asymmetric junctions between a presynaptic neuron and a postsynaptic target with different molecular composition, structure, and activities. Bruchpilot (Brp), Liprin alpha (Lip α) and Synaptotagmin 1 (Syt1) are conserved proteins localized in the presynaptic side.

Brp is a well studied component of the presynaptic component of the synapses in *Drosophila* that accumulates in mature active zones (AZ). Brp is the orthologue of human AZ protein ELKS/CAST/ERC, and it is required for synapse formation (Wagh et al., 2006). Lip α is a presynaptic scaffolding protein, orthologue to several human genes including PPFIA1 (PTPRF interacting protein alpha 1) and PPFIA2 (PTPRF interacting protein alpha 2). Lip α directly interacts with tyrosine phosphatase receptors and it is involved in synapse formation, anterograde synaptic vesicle transport, neuron development, synapse organization and axon guidance (Astigarraga et al., 2010; Fouquet et al., 2009; Kaufmann et al., 2002). Finally, Syt 1 is a pre-synaptic vesicle calcium binding protein that functions as the fast calcium sensor for neurotransmitter release at synapses (Yoshihara and Montana, 2016).

Synapses elicit neurotransmission by mediating the clustering and fusion to the plasma membrane of neurotransmitter containing vesicles which release into the synaptic space (Chou et al., 2020). The postsynaptic side is characterized by the accumulation of neurotransmitter receptors, including Glutamate receptors (GluR), the protein discs large (Dlg), orthologue of human PSD95 protein which mediates the clustering of postsynaptic molecules (Koh et al., 1999), and Synaptotagmin 4 (Syt 4), a vesicular calcium binding protein, directly implicated in retrograde signaling at synapses. Syt 4 is proposed to regulate calcium-dependent cargo trafficking within the postsynaptic compartment (Harris et al., 2016).

Benefitting from the conserved nature of most synaptic components, we set out to dissect the pre-versus post-synaptic contributions to GB progression using a Drosophila model of the human disease, in which the pathological condition of each cell type can be genetically manipulated. Thus, in addition to demonstrate that neuron-glioblastoma synaptogenesis is a conserved mechanism in GB progression, we show that synapses are also formed intratumoral and identify several synaptic genes required for GB expansion and premature death.

## MATERIALS AND METHODS

### Fly Stocks

Flies were raised in standard fly food at 25ºC, otherwise indicated.

Fly stocks used were *UAS-lacZ* (BL8529), *UAS-myr-RFP* (BL7119), repo-Gal4 (BL7415), *tub-gal80ts* (BL7019), *elav-lexA* (BL 52676), *UAS-CD2::HRP* (BL8763), *UAS-Syt1-GFP, lexAop-nSyb-spGFP1-10UAS-CD4-spGFP11* (BL64315), *UAS-nSyb-spGFP1-10lexAop-CD4-spGFP11* (BL64314), *UAS-mLexA-VP16-NFAT lexAop-rCD2-GFP* (CaLexA, BL66542), *UAS-Syb*^*RNAi*^ *(BL38234), UAS-Liprin-alpha*^*RNAi*^*(BL53868), UAS-Syt1*^*RNAi*^*(BL31289), UAS-Syt4*^*RNAi*^*(BL39016), UAS-Brp*^*RNAi*^*(BL25891)* from the Bloomington Stock Center (https://bdsc.indiana.edu/index.html), *UAS-yellowRNAi* (KK106068), *UAS-GluRIIA-RNAi* (KK101686), *UAS-Dlg-RNAi* (KK109274), *UAS-Bruchpilot-RNAi* (KK104630) from the Vienna Drosophila Resource Centre (https://stockcenter.vdrc.at/control/main), *UAS-dEGFR*λ, *UAS-dp110CAAX* (A gift from R. Read), UAS-Liprinα-GFP, *GluRIIA-GFP*, (A gift froom S.J. Sigrist).

### Inmunohistochemistry

Third-instar larval brains, were dissected in phosphate-buffered saline (PBS), fixed in 4% formaldehyde for 30min, washed in PBS + 0.1 or 0.3% Triton X-100 (PBT), and blocked in PBT + 5% BSA for 1 hour. Samples were incubated overnight with primary antibodies diluted in block solution, washed in, incubated with secondary antibodies diluted in block solution for 2 hours and washed in PBT. Fluorescent labeled samples were mounted in Vectashield mounting media with DAPI (Vector Laboratories).

Primary antibodies used were: mouse anti-Repo (DSHB 1:200), rabbit anti-GFP (Invitrogen A11122, 1:500), mouse anti-GFP (Invitrogen A11120, 1:500), mouse anti-Nc82(brp) (DSHB 1:30), rabbit anti-GluRIID (1:100) (gift from Dr. Stephan Sigrist, European Neuroscience Institute, Göttingen, Germany, mouse anti-ELAV (DSHB 1:50), Rabbit anti-Hrp (Jackson Immunoresearch 111-035-144, 1:400).

Secondary antibodies used were: anti-mouse Alexa 488, 568, 647, anti-rabbit Alexa 488, 568, 647 (Thermofisher, 1:500).

Samples were analyzed by confocal microscopy (LEICA TCS SP5).

### TEM

Transmission electron microscopy (TEM) was performed in CNS of 3rd instar larvae with horseradish peroxidase (HRP) genetically driven to glial cells (*repo-Gal4:UAS-HRP CD2*). Brains were fixed in 4% formaldehyde in PBS for 30 min at room temperature, and washed in PBS, followed by an amplification of HRP signal using the ABC kit (Vector Laboratories) at room temperature. After developing with DAB, brains were washed with PBS and fixed with 2% glutaraldehyde, 4% formaldehyde in PBS for 1h at room temperature. After washing in a phosphate buffer the samples were postfixed with OsO4 1% in 0.1 M 7phosphate buffer, 1% K3[Fe(CN)6] 1h at 4ºC. After washing in dH2O, Brains were incubated with tannic acid in PBS for 1min at room temperature then washed in PBS for 5min and dH2O 2×5min. Then the samples were stained with 2% uranyl acetate in H2O for 1h at room temperature in darkness followed by 3 washes in H2O2d. Brains dehydrated in ethanol series (30%, 50%, 70%, 95%, 3×100% 10 min each at 4ºC). Infiltration: samples were incubated in EtOH:propylene’s OXID (1:1;V.V) for 5 min, propylene’s OXID 2×10min, propylene’s OXID:Epon (1:1) for 45 min, Epon 100% in agitation for 1 h and Epon 100% in agitation overnight. Then change to Epon 100% for 2-3 h. After, the samples were encapsulated in BEEM and incubated 48h at 60ºC for polymerization. Finally, the samples were cut in ultra-fine slices for TEM imaging (Martín-Peña et al., 2014).

### Imaging

Fluorescent images were acquired by confocal microscopy (LEICA TCS SP5) and were processed using Fiji (Image J 1.50e). These images were quantified with Fiji (Image J 1.50e) or Imaris 6.3.1 (Bitplane) software. The images of the ultra-fine slices were taken with a Transmission electron microscopy JEM1010 (Jeol) with a CMOS TemCam F416 (TVIPS) camera and processed with Adobe Photoshop CS4. Figures were assembled using Adobe Photoshop CS4 and Adobe Illustrator CS4.

IMARIS quantification (Imaris 6.3.1 software): The number of glial cells (Repo+) and the number of synaptic active sites was quantified by using the spots tool. The tumor volume was quantified using the surface tool. We selected a minimum size and threshold for the punctae or surface in the control samples of each experiment to establish the conditions. Then we applied the same conditions to the analysis of each corresponding experimental sample.

#### Fiji quantification

##### - CaLexA expression

We used the NFAT-based neural tracing method-CaLexA (calcium-dependent nuclear import of LexA)-for labeling active neurons in behaving animals. CaLexA (green) signal intensity where determined using ImageJ to calculate the mean grey value of each brain lobe.

##### - Syt1-GFP expression

Syt1-GFP (green) signal and glia membrane myrRFP (red) signal intensity were determined using ImageJ (mean grey value) in three single slices at the middle of each brain lobe to calculate the ratio GFP/RFP.

All samples were treated, acquired and measured under the same conditions and in parallel

##### - GRASP

We used a modified version of this system to specifically detect synaptic contacts. It is based on the fusion of synaptobrevin protein (Syb) to the 1-10 fragment of GFP (Syb-GFP_1-10_), and the expression of a membrane bound form of the 11 fragment of GFP (CD4-GFP_11_). UAS-nSyb*-spGFP*^*1-10*^ : lexAop-*CD4-sp GFP*^*11*^ (BL62314) and lexAop-*nSyb-spGFP*^*1-10*^ : UAS-*CD4-sp GFP*^*11*^ (BL62315) were expressed in neurons (*elav-lexA*) and glial (*repo-Gal4*) cells respectively. These complementary GFP fragments will reconstitute a functional fluorescent reporter at the points of contact and therefore, it will allow the identification of the presynaptic and postsynaptic cells (e.g. glia and neuron) (Karuppudurai et al., 2014).

### Viability assays

Flies were crossed at restricted temperature (17 °C, to inactivate the *UAS/Gal4* system with *tub-Gal80ts*) for 4 days then progeny was transfer at 29°C (when the *UAS/Gal4* system is active and the glioblastoma develops). The number of adult flies emerged from the pupae were counted for each genotype. The number of control flies was considered 100% viability and all genotypes are represented relative to controls. Experiments were performed in triplicates.

### Survival assay

For survival analyses of adult flies, males and females were analyzed separately. 0-5 day old adult flies raised at restricted temperature were put at 29°C in groups of 10 animals per vial and were monitored blinded every 2-3 days; each experiment was done at least three times.

### Quantifications and Statistical Analysis

All experiments including different genotypes were done in parallel under the same experimental conditions, with the exception of viability analysis where each genotype was normalized with their parallel control. Data were analysed and plotted using GraphPad Prism v7.0.0 and Excel (viability assays). A D’Agostino & Pearson normality tests were performed and data with normal distributions were analysed using a two-tailed T-test with Welch-correction. If data had multiple comparisons, a One-way ANOVA with Bonferroni posthoc-test was used. Data that did not pass normality testing were submitted to a two-tailed Mann-Whitney U-test or where the data had multiple comparisons a Kruskal-Wallis test and Dunnett’s post hoc-test. Error bars represent Standard Error of the Mean, significance values are: ***p≤0.0001, ** p≤0.001, *p≤0.005, ns=non-significant.

## Acknowledgements

We thank Professor Alberto Ferrús and Dr. Paco Martín for critiques of the manuscript and for helpful discussions. Esther Seco for fly stocks maintenance. We want to thank the Vienna *Drosophila* Resource Centre, the Bloomington *Drosophila* stock Centre and the Developmental Studies Hybridoma Bank for supplying fly stocks and antibodies, and FlyBase for its wealth of information. We acknowledge the support of the Confocal Microscopy unit and Molecular Biology unit at the Cajal Institute and the *Drosophila* Transgenesis Unit and the Transmission Electron Microscope unit at CBMSO for their help with this project. Research has been funded by grant PID2019-110116GB-100 from the Spanish Ministerio de Ciencia e Innovación and a Postdoctoral Fellowship from the Comunidad de Madrid (2016-T2-BMD-1295 to M.L.-P.). Authors declare no conflicts of interest.

## Results

We performed a *Drosophila* biased genetic screening to search for relevant genes related to GB progression. We selected 2000 genes involved in cell to cell communication, and we used VDRC UAS-RNAi lines to knockdown the expression of such genes encoding transmembrane, secreted and cell to cell communication proteins. In addition, we used the previously validated EGFR/PI3K model (Portela and Casas-Tintó, 2020; Portela et al., 2019b, 2020; Read et al., 2009). GB induction in larvae causes premature death and animals do not reach adulthood. We took advantage of this unequivocal phenotype as a read-out, quantifying the number of adult flies that emerged from each experiment. We obtained 25 RNAi lines that rescued the lethality caused by the GB. Among the suppressors, we found well known mediators of GB progression such as *Frizzled1* (*Fz1*) or *Gryzun (Gry)* receptors and PI3K signaling pathway members (Portela et al., 2019b, 2019a, 2020). These genes validate the experiment as positive controls. Most RNAi lines, as well as negative controls UAS-*yellow* RNAi or UAS-*beta-galactosidase (lacZ)*, did not rescue GB-induced pupal lethality however, we found genes encoding synaptic proteins, such as *liprin* α *(lip* α*) and synaptotagmin1 (Syt 1)* that rescue GB-induced lethality (Figure 1A). These results motivated this study to determine the contribution of synaptic components to GB progression.

**Figure 1.**
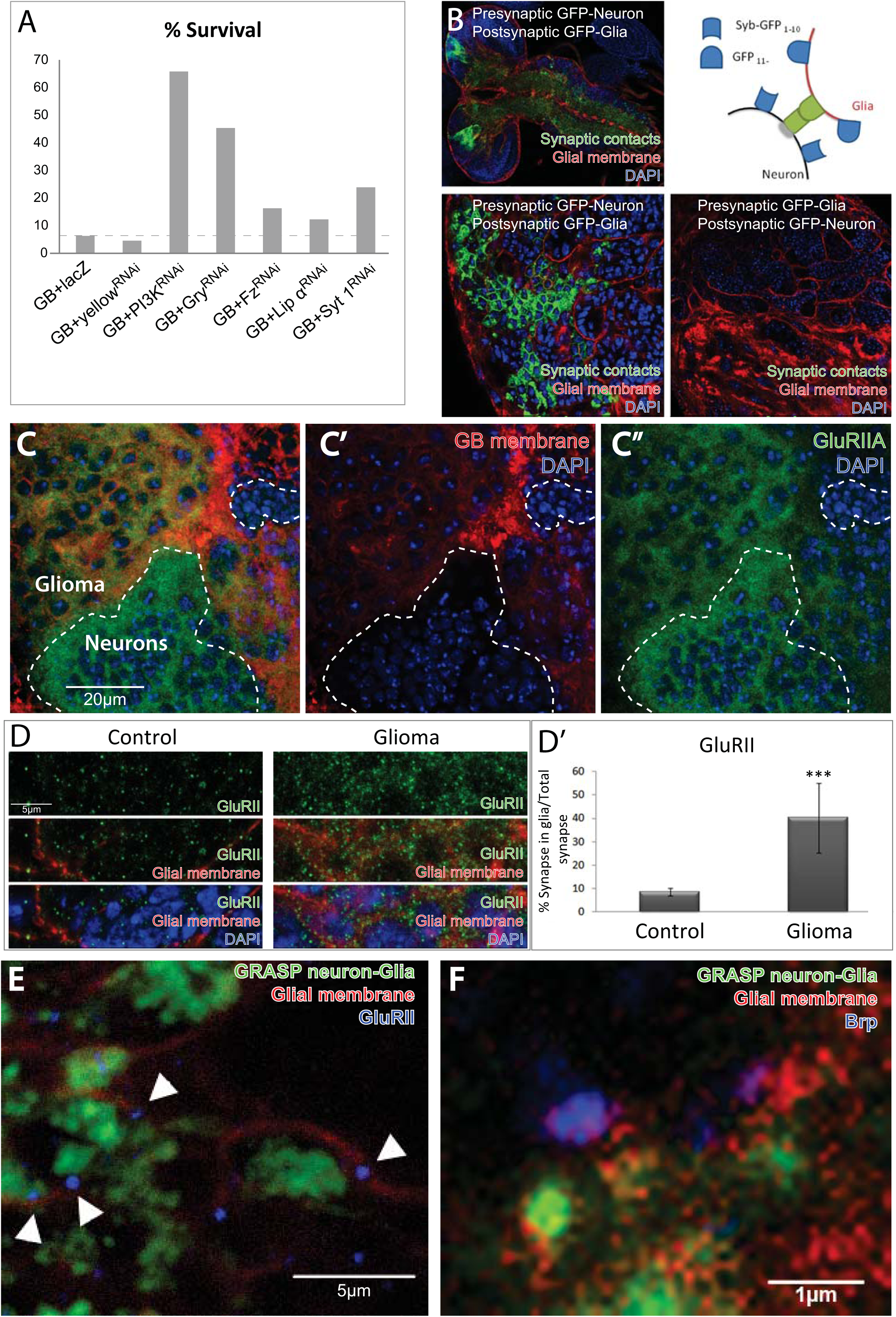
GB cells form synapses with neurons. A) Histogram showing the percentage of viability of flies when glioblastoma (GB) is induced alone (GB+lacZ or GB+yellowRNAi) or combined with *PI3K, gry, fz, lip* α or *syt 1* knockdown in GB cells by RNAi. Percentage corresponds to the number of adult flies that emerged from the pupae for each genotype, relative to the controls (siblings without *repoGal4*, considered 100% of viability). B top left) confocal image of GRASP+ signal in larval brain when presynaptic Syb-GFP_1-10_ fragment is expressed in neurons and CD4-GFP_11_ fragment is expressed in GB cells; B bottom left) magnification of above; B top right) Diagram of GRASP technique; B bottom right) confocal image of GRASP – signal when presynaptic Syb-GFP_1-10_ fragment is expressed in glioma cells and CD4-GFP_11_ fragment is expressed in neurons; Synaptic contracts are shown in green (GFP), glial membrane are in red (repo>myrRFP) and all nuclei (DAPI) are in blue. C) Confocal image of larval GB brain showing glial membrane (red) surrounding healthy tissue (neurons, not red) and stained with anti-GluRIIA (green). Dotted lines mark the limit between neurons and glioma cells. GluRIIA postsynaptic protein is detected in both GB and healthy tissue (C and C’’). DAPI staining nuclei is in blue. D) GluRIID signal (green) in control and GB larval brains, presented with glial membrane or glial membrane and DAPI in the bottom images. D’) Number of GluRIID-positive dots overlapping with glial membranes in control and GB samples. Statistic: Unpaired T-Test. E) High magnification confocal image showing GB membrane (red) and Neuron-Glia GRASP signal (green) in the proximity of Brp signal (blue). F) High magnification confocal image showing GB membrane (red) and Neuron-Glia GRASP signal (green) in the proximity of GluRIID signal (blue). Scale bars: 20μm (C), 5μm (D and E) and 1μm (F).

### Neurons produce synaptic contacts with glioma cells

It was recently described that neurons establish functional synapses with glioblastoma cells in mouse xenografts (Venkataramani et al., 2019; Venkatesh et al., 2019). In these studies GB cells are postsynaptic, however our results from the screening indicate that presynaptic proteins are also involved in GB-induced lethality (Figure 1A). Therefore, we wondered if GB cells were pre- or postsynaptic in the *Drosophila* GB model. We used the GFP reconstitution across synaptic partners (GRASP) technique (Macpherson et al., 2015) to determine synaptic contacts between GB cells and neurons. This technique allows the identification of pre- and postsynaptic cells. The confocal images of larvae brains show that GFP signal is reconstituted (GRASP+) if presynaptic Syb-GFP_1-10_ fragment is expressed in neurons, under the control of the specific neuronal enhancer *elav-lexA*, and CD4-GFP_11_ fragment is expressed in GB cells under the control of the specific glial enhancer *repo-Gal4* (Casas-Tintó et al., 2017) (Figure 1B). In addition, we co-expressed a myristoylated form of Red Fluorescent Protein (myrRFP) in glial cells under the control of the UAS/Gal4 system to visualize GB cells membranes. In contrast, GFP does not reconstitute when GB cells express the presynaptic component of GRASP, and neurons express the post-synaptic component (Figure 1B). These results indicate that not tumoral healthy neurons (pre-synaptic) establish synapses with GB cells (post-synaptic) in *Drosophila*. These contacts occur in a unidirectional manner, therefore validating previous results in other GB model systems.

Next, to further explore the postsynaptic role of GB cells, we studied the expression of the post-synaptic *Glutamate receptor II* (*GluRII)* gene in GB cells. We used a GluRII protein trap transgenic line to monitor the expression and localization of GluRIIA-GFP protein. Confocal images (Figure 1C) show GB tissue (red) and not-tumoral healthy tissue (not-red), and we observed the presence of GluRIIA-GFP signal in GB tissue (Figure 1C’). Moreover, to confirm the presence of GluRII protein in the membranes of GB cells, we used a validated antibody against the GluRIID subunit. Confocal images of control larval brain samples showed GluRIID dotted signals through the brain revealing glutamatergic synapses (Figure 1 D). We quantified the number of GluRIID positive dots that overlap with glial membranes in control and GB samples. The results show that less than 10% of the total GluRIID signal corresponds to glial membranes in control samples, whereas the number of GluRIID positive signals in the GB membrane reaches 40% of total synapses (Figure 1D-D’). In summary, our data suggest that GluRIID protein accumulates in GB cells.

To further determine the nature of these synaptic structures, we co-stained GB larval samples with a specific monoclonal antibody that recognizes the pre-synaptic protein Bruchpilot (Brp), or an antibody against GluRIID and analyzed the relative position with GRASP signal. High magnification confocal images show GB membrane (red) and Neuron-Glia GRASP signal (green) in the proximity of Brp signal (blue in Figure 1 E) or GluRIID (blue in Figure 1 F). In both cases, proteins appear at less than 1 micrometer distance (Figure1 E-F) compatible with the formation of synapses. These results suggest that pre- and post-synaptic proteins accumulate in the GB-neuron contact region.

### GluRIIA and dlg post-synaptic proteins are required for GB expansion

Once we have demonstrated the postsynaptic nature of GB cells we aimed to determine their contribution to GB progression. Third instar larvae brains with GB display expanded glial membrane, formation of perineuronal nests (Casas-Tintó and Portela, 2019) (magenta in Figure 2 A) and a subsequent increase of brain volume (Figure 2A). We used RNAi techniques to knockdown *dlg* or *GluRII* postsynaptic genes in GB cells. The quantifications of the results show that *dlg* or *GluRII RNAi* prevents membrane expansion, disrupts perineuronal nest formation and prevents brain size increase (Figure 2B-B’ and C).

**Figure 2.**
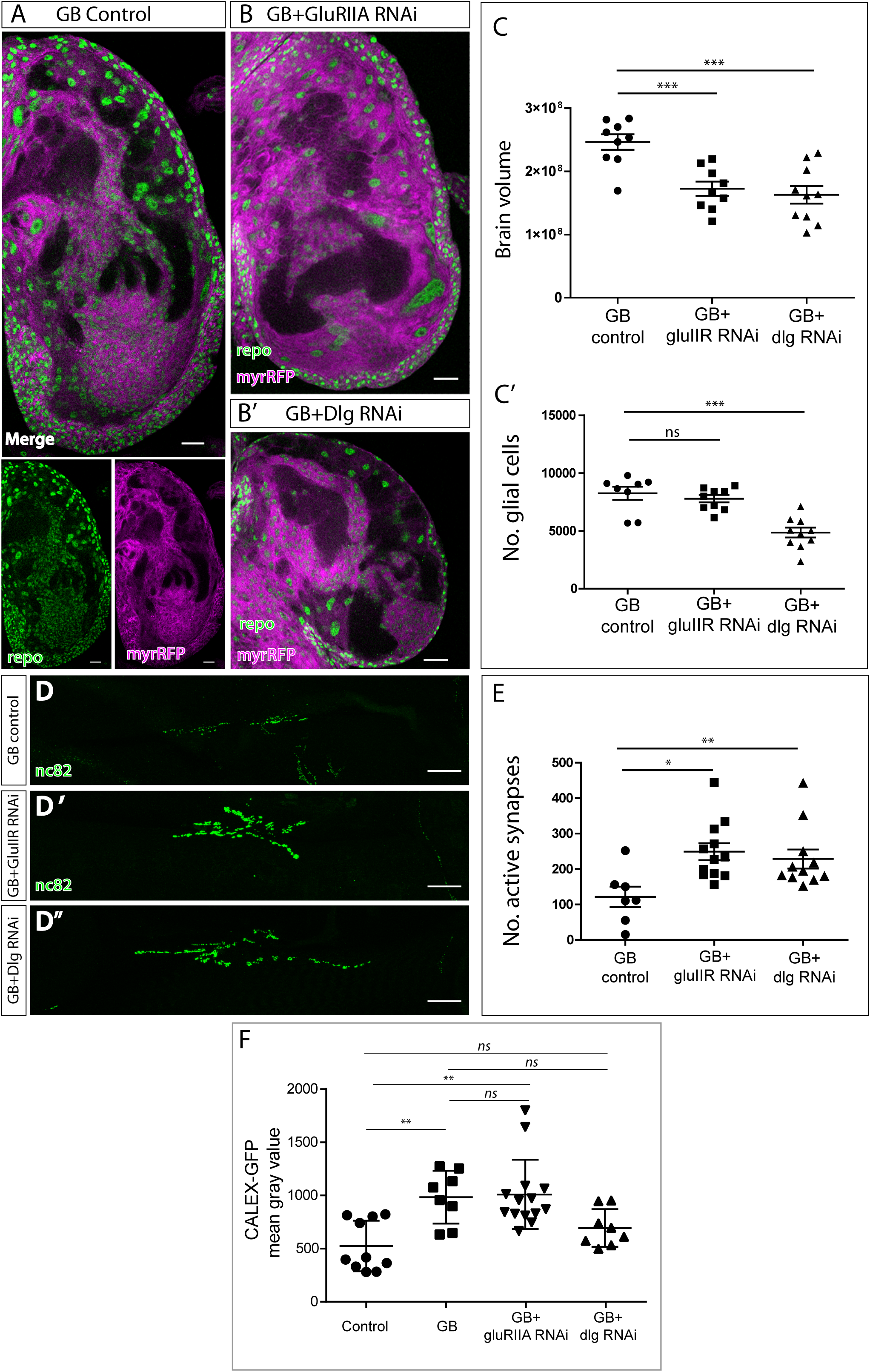
Postsynaptic proteins GluRII and Dlg are required for GB progression. A) Representative confocal image of GB larval brain lobe showing glial nuclei (green) and glial membrane (magenta). Bottom images show green and magenta channels separately. B-B’) Representative confocal images of GB larval brain lobe + *GluRIIA RNAi* expression (B) or *expression* (B’) in GB cells. Glial nuclei are in green and glial membrane in magenta. C) Quantification of glial membrane volume (C) or number of glial cells (C’) in GB control and GB with *GluRIIA* or *dlg* RNAi. Statistics: Dunnett’s Multiple Comparison Test. D) Confocal images of Neuromuscular junctions showing Brp accumulation in GB controls (D) and GB with *GluRIIA RNAi* (D’) or *dlg* RNAi (D’’). E) Quantification of synapse number (Brp positive dots) in GB controls and GB with *GluRIIA RNAi* or *dlg RNAi*. Statistics: Dunnett’s Multiple Comparison Test. E) Quantifications of CALEX signal control brains, GB brains and GB upon postsynaptic genes *dlg* or *GluRIIA* knockdown. Statistics: Dunnett’s Multiple Comparison Test. Scale bars: 50 μm (A, B and D-D’’)

Next, we analyzed the contribution of *dlg* and *GluRII* to the increase of glial cell number in GB, we stained brain samples with the specific anti-repo antibody to visualize glial nuclei (Figure 2B-B’). The quantifications show that *dlg* knockdown reduces significantly the number of GB cells, but, on the contrary, *GluRIIA RNAi* expression does not prevent the increase in glial cell number in GB (Figure 2 C’).

Given that downregulation of *GluRII* or *dlg* prevent the GB membrane expansion (Figure 2 C) and that the expansion of tumor microtubes in GB cells mediates the reduction of synapses in neighboring healthy neurons (Jarabo et al., 2021; Portela et al., 2019b, 2020), we wondered if *GluRII* and *dlg* were required for the GB-induced neurodegeneration. We analyzed larval neuromuscular junctions (MNJ) which is a validated system to analyze neurodegeneration (Arnés et al., 2020; Jordán-Álvarez et al., 2017). We quantified the number of synapses by counting the number of active zones (Brp positive dots, green in Figure 2 D-D’’). The results indicate that GB progression causes a reduction of synapse number, as previously reported, and that *GluRII* or *dlg* knockdown in GB cells prevents the reduction of synapse number in neurons, preventing neurodegeneration (Figure 2 F). Thus, the expression of *dlg* and *GluRII* post-synaptic genes is required for GB-induced neurodegeneration.

Cell proliferation in GB is associated with calcium-mediated activity (Venkataramani et al., 2019), thus we analyzed the contribution of *dlg* or *GluRIIA* to calcium activity in GB cells. To monitor calcium activity, we used the CaLexA system (see Materials and Methods). Quantification of CaLexA signal showed a significant increase of calcium signal in GB samples, but *dlg* knockdown in GB cells maintained calcium levels as controls (Figure 2 G). However, we did not find significant differences upon *GluRIIA* downregulation. These results indicate that GB cells show enhanced calcium-dependent activity, in line with previous data from other GB models (Venkatesh et al., 2019). Moreover, our data indicate that this enhanced calcium activity is dependent on *dlg* expression, while independent on that of *GluRIIA*.

### Vesicle calcium binding proteins are required for GB progression

Vesicle calcium binding, vesicle transport and neurotransmitter release are cellular mechanisms related to synaptic function (Kavalali, 2015; Südhof, 2013). We have found that GB has an enhanced calcium activity that can be reduced by downregulating the expression of *dlg*. Moreover, our screening results indicate that downregulation of *Syt 1*, which encodes a presynaptic Calcium-binding protein, partially rescued the lethality caused by the GB (Figure 1 A). This motivated the study of Synaptotagmin 1 (Syt 1), as well as Synaptotagmin 4 (Syt 4, a postsynaptic Ca-binding protein) in GB progression.

To explore the pre-synaptic role of GB, we analyzed Syt 1 accumulation in normal glia and GB cells. We used a Syt1-GFP fusion protein to monitor *Syt 1* expression and measure the accumulation of Syt 1 in glial membranes, comparing normal glia with GB cells. The confocal images show that Syt 1 accumulates in the membrane of glial cells, and this accumulation exacerbates in GB samples (Figure 3 E-F). This result indicates that components from the presynaptic side also contribute to GB progression.

**Figure 3.**
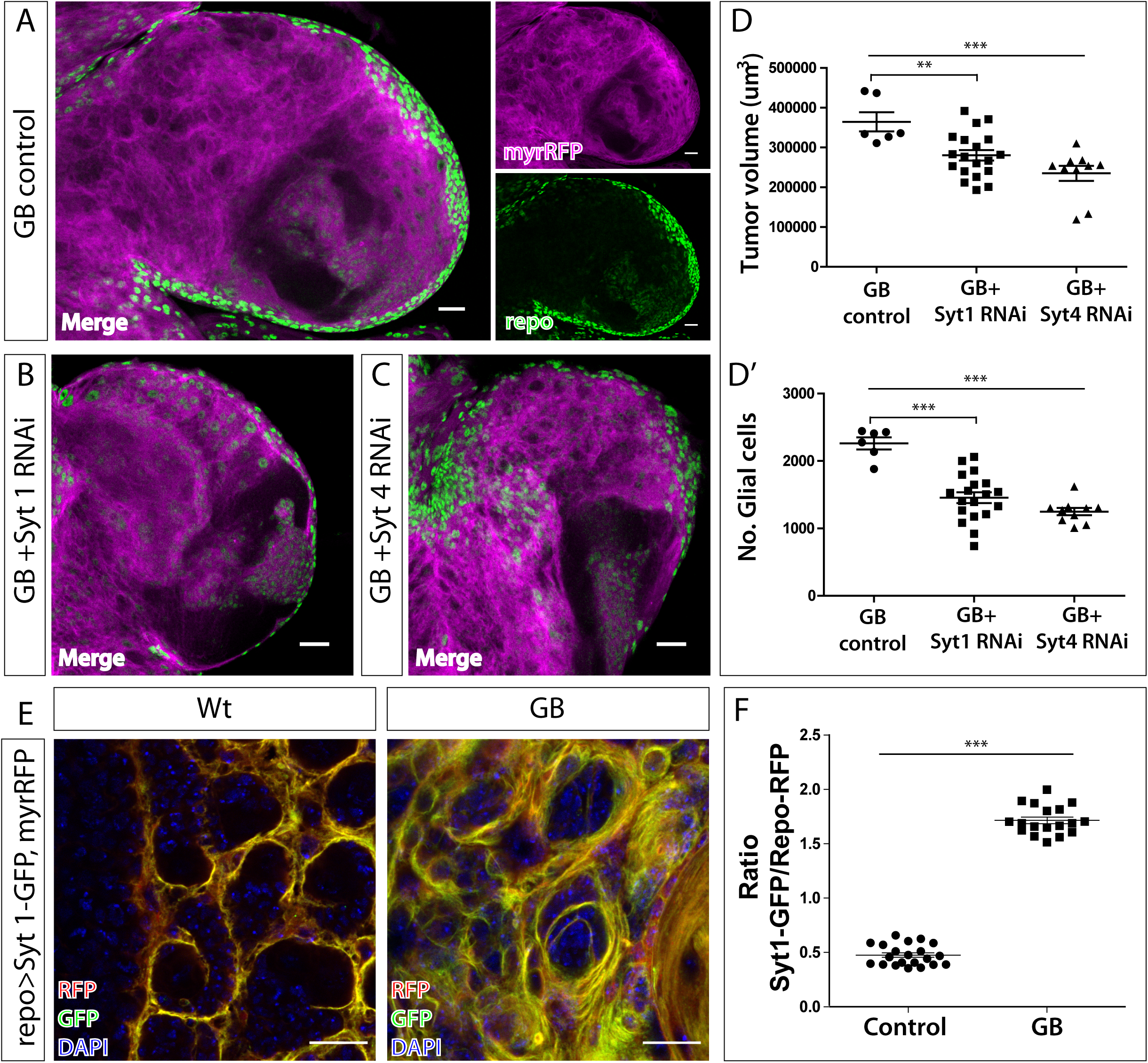
Sygnaptotagmin 1 and 4 are required for GB progression. A) Representative confocal image of GB larval brain lobe showing glial nuclei (green) and glial membrane (magenta). Right images show green and magenta channels separately. B-C) Representative confocal images of GB larval brain lobe expressing *Syt 1 RNAi* (B) or *Syt 4 RNAi* (C). Glial nuclei are shown in green and glial membranes in magenta. D) Quantification of glial membrane volume (D) or number of glial cells (D’) in GB control and GB + *Syt 1* or *Syt 4* knockdown. Statistics: Dunnett’s Multiple Comparison Test. E) Representative confocal images showing Syt 1-GFP accumulation (green) in control (left) and GB brains (right). Glial membrane is in red and nuclei (DAPI) in blue. F) Quantification of GFP/RFP ratio, corresponding to Syt1-GFP/glial membrane in control and GB brains. Statistics: Unpaired T-Test with Welch’s correction. Scale bars: 50 μm (A-C) and 15 μm (E)

We used specific RNAi tools to knockdown *Syt 1* or *Syt 4* expression, and studied the effects on glial cell number and GB volume. The quantifications showed that *Syt 1* or *Syt 4* knockdown specifically in GB cells prevents the expansion of GB and the over proliferation of GB cells (Figure 3A-D’). Therefore, the expression of these two genes that regulate vesicle transport and neurotransmitter release are required for GB development in *Drosophila*. These results support the post-synaptic nature of GB cells (*Syt 4*), and also support a pre-synaptic condition (*Syt* 1) of GB cells as a requirement for GB expansion.

### Presynaptic proteins are required for the enhanced GB calcium activity

To further explore the presynaptic condition of GB cells, we measured the accumulation of intracellular calcium by knocking down presynaptic genes such as *Brp, Syt1* or *Lip*α in GB cells. The quantification of CaLexA intensity signal showed that the knockdown of these presynaptic genes prevented the increase of CaLexA signal in GB cells (Figure 4 A). This suggests that presynaptic genes are required for the induction of calcium accumulation and calcium-dependent activity in GB cells.

**Figure 4.**
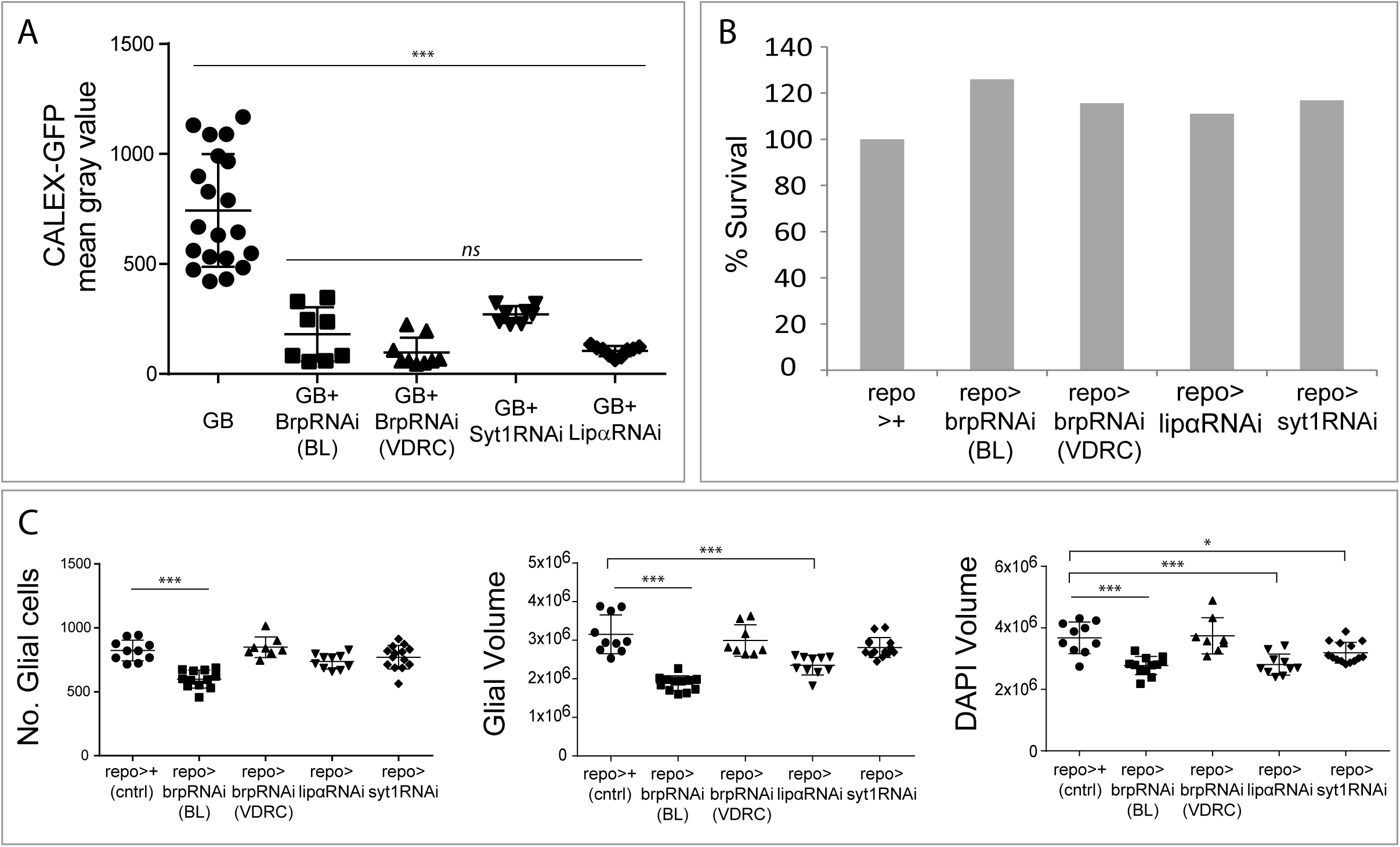
Presynaptic proteins are required for Calcium influx in GB. A) Quantifications of CaLexA signal in GB brains upon presynaptic genes brp, Syt 1 or lip □ knockdown by RNAi. Statistics: Bonferroni’s Multiple Comparison Test. B) Histogram showing the viability of control flies compared with flies where *brp, lip* α or *syt 1* are downregulated in glia. C) Quantification of the number of glial cells, glial membrane volume and DAPI volume, per larval brain lobe in wt controls, and animals where *brp, lip* α or *Syt 1* are downregulated in glia. Statistics: Dunnett’s Multiple Comparison Test.

To ensure that presynaptic proteins have a specific role in GB progression and are not required for normal glia development we analyzed the viability of animals where *lip* α, *brp* or *Syt 1* are knocked down in glial cells (through all developmental stages under the control of *repo-Gal4)*. The quantification shows that, in all cases, the percentage of animals that reach adulthood compared with controls (the siblings) is similar or even higher (Figure 4 B), and therefore, we conclude that downregulation of *lip* α, *brp* or *Syt 1* in glial cells does not affect viability and therefore, are not required for vital functions during development.

Additionally, we dissected brains of third instar larvae and quantified the number of glial cells, the volume of glial membrane and brain size (Figure 4 C). The results show that the number of glial cells was normal in all cases with the exception of the expression of *Brp RNAi BL*. In addition, we measured the total volume of glial cells membrane marked with myrRFP. The quantification of RFP volume indicates that the expression of *Brp RNAi VDRC* and *Syt 1 RNAi* did not cause any effect during development; however *Brp RNAi BL* and *Lip* α *RNAi* expression in glial cells caused a reduction of the total volume of glial cells membranes (Figure 4 C). Finally, we measured the total brain volume marked with DAPI, brain volume was reduced upon *Brp VDRC RNAi, Lip* α *RNAi* and *Syt 1 RNAi* expression, but not upon *Brp RNAi BL* expression (Figure 4 C). These data unveil a role of these synaptic proteins in the biology of normal glial cells that was unknown hereto.

### Intratumoral synapses in GB

The results obtained so far indicate the presence of presynaptic proteins in GB cells. Moreover, the presynaptic nature of GB cells has been demonstrated by the requirement of presynaptic proteins to calcium signal enhancement. Does that imply synaptogenesis between GB cells? We found Syt 1 accumulation in GB compared to controls (Figure 3 E-F) concomitant with the lethality rescue observed upon S*yt 1* downregulation in GB (Figure 1A). *Lip* α downregulation in GB cells also rescues lethality (Figure 1A), as well as calcium enhancement (Figure 4 A), besides, to further confirm the presence of Lip α in GB we quantified Lip α accumulation in normal glia and GB cells using a Lip-GFP reporter line. The data show accumulation of Lip α -GFP in dots at the membrane of glial cells, and this accumulation increases in GB samples (Figure 5 A-A’). This result shows that the localization and accumulation of the pre-synaptic protein Lip α is enhanced in GB cells, compatible with the presynaptic nature of GB cells and the formation of synapses. By contrast, however, GRASP experiments had suggested that GB cells only function as post-synaptic structures with respect to neurons (see above, Figure 1B).

**Figure 5.**
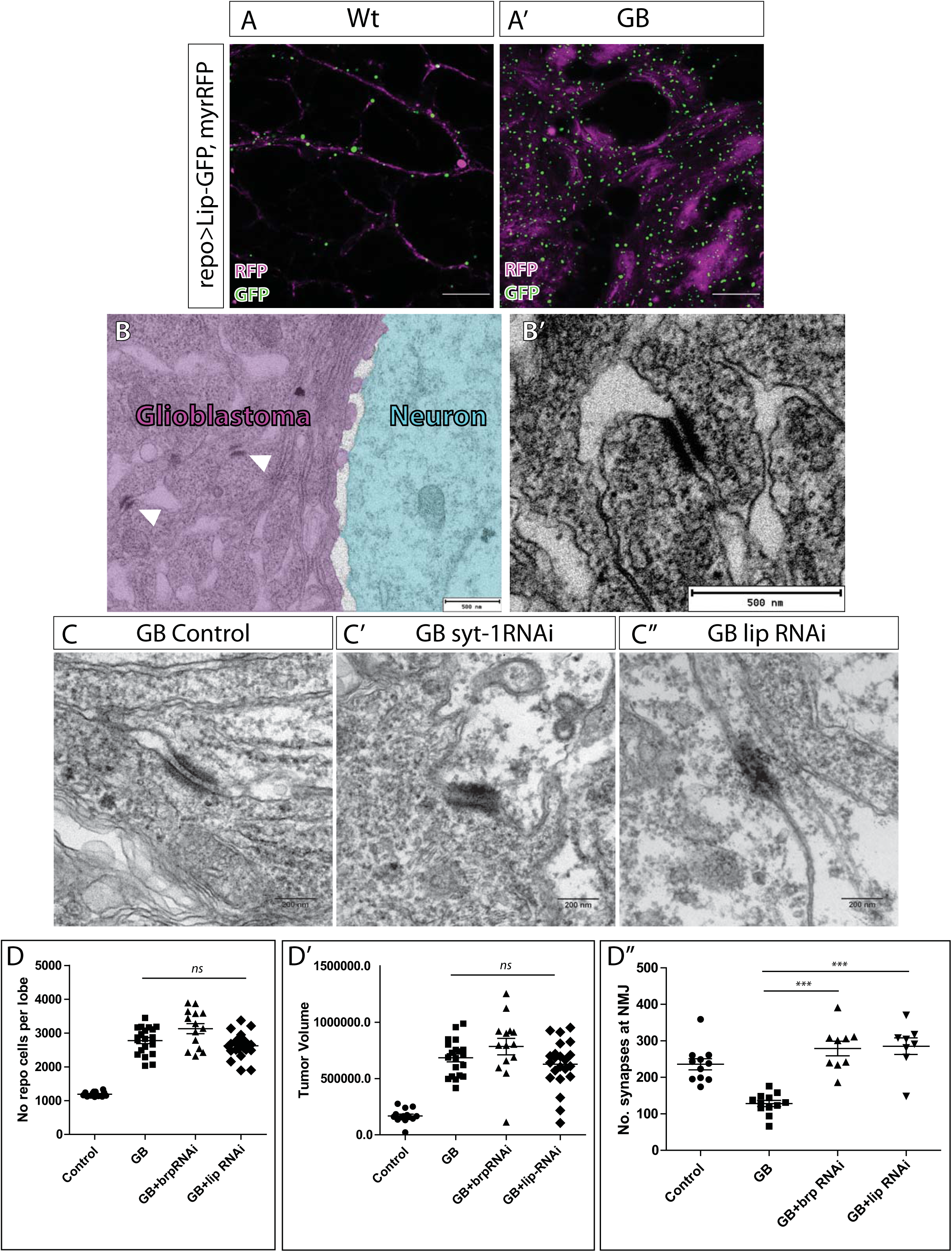
GB intratumoral synapses. A) Representative confocal images of Lipα-GFP accumulation in glial cells in control (A) and GB (A’) samples. Lip-GFP is in green and GB membrane in magenta. B) Representative TEM images of GB samples. B) GB is colored in magenta and neurons are in cyan. White arrowheads indicate electron dense signals between GB named “Intratumoral synapses”. B’) TEM Magnification of the intratumoral synapses. C) Comparison of Intratumoral synapses in GB (C) and GB + *Syt 1 RNAi* (C’) or *lip* α *RNAi* (C’’) expression. D) Quantification of glioma cells number (D), tumor volume (D’) or synapses number at the NMJ (D’’) in control wt, GB, GB with Brp knocked down or GB with lip-α knocked down samples. Statistics: One-way ANOVA (D-D’’) and Bonferroni’s Multiple Comparison Test (D’’). Scale bar: 15 μm (A-A’), 500 nm (B-B’), 200 nm (C-C’’).

To clarify if pre-synaptic proteins indeed have a role in GB progression, we explored the possibility of synapses formation within the tumoral mass between GB cells, here defined as “intratumoral synapses”. To that end, we marked GB membranes with HRP and obtained transmission electron microscopy (TEM) images of GB samples and controls. EM images show high density structures between GB cells that are compatible with synapses (Figure 5B-B’). To validate if these densities exhibit synaptic features, we knocked down *Syt 1* or *Lip* α in GB cells and analyzed the tissue under TEM. The images show a morphological disruption of these electron densities upon *Syt 1* knockdown (Figure 5C-D) as well as upon *lip* α knockdown (Figure 5 C and E). These results suggest that the pre-synaptic proteins Syt 1 and Lip α are functionally required to build these putative GB-GB Intratumoral synapses.

Next, we analyzed the contribution of Brp and Lip α pre-synaptic proteins to GB progression. We quantified the number of GB cells and the volume of the tumoral mass upon *brp* or *lip* α knockdown selectively in GB cells. The quantification of confocal images shows that *brp* or *lip* α *RNAi* do not reduce the number of GB cells, nor the volume of the tumor (Figure 5F-G). Nevertheless, our screen results indicate that Lip α is required for GB causing lethality (Figure 1 A), thus, we investigate the requirement of Brp and Lip α for GB neurodegeneration. We quantified the number of synapses at the NMJ in GB control larvae and GB knocking down *lip* α or *brp* specifically in GB cells. The results show that downregulation of *brp* or *lip* α rescues synapse number at NMJ to normal values (Figure 5H) and therefore prevents neurodegeneration caused by GB. Thus, the expression of *brp* or *lip* α in glia is required to cause the GB characteristic synapse number reduction and neurodegeneration,

### Synaptic genes rescue premature death caused by GB

Finally, to determine the systemic impact of synaptic genes in GB, we evaluated the contribution of synaptic genes to the premature death caused by GB progression (Portela et al., 2019b, 2019a). We measured the life span of adult male and female adult flies with GB, and compared it with flies in which certain pre- or post-synaptic genes had been knocked-down selectively in GB cells. The survival curves show in all cases that the induction of GB reduces the survival of male and female adult flies (Figure 6, grey lines). However, the knockdown of *brp* or *lip* α prevents GB lethality and fully restores life span to control levels in males and females (Figure 6 A). Also, the knockdown of *Syt1* also prevents GB-induced lethality (Figure 6 B) suggesting that the expression of these pre-synaptic genes is required in GB cells to cause premature death.

**Figure 6.**
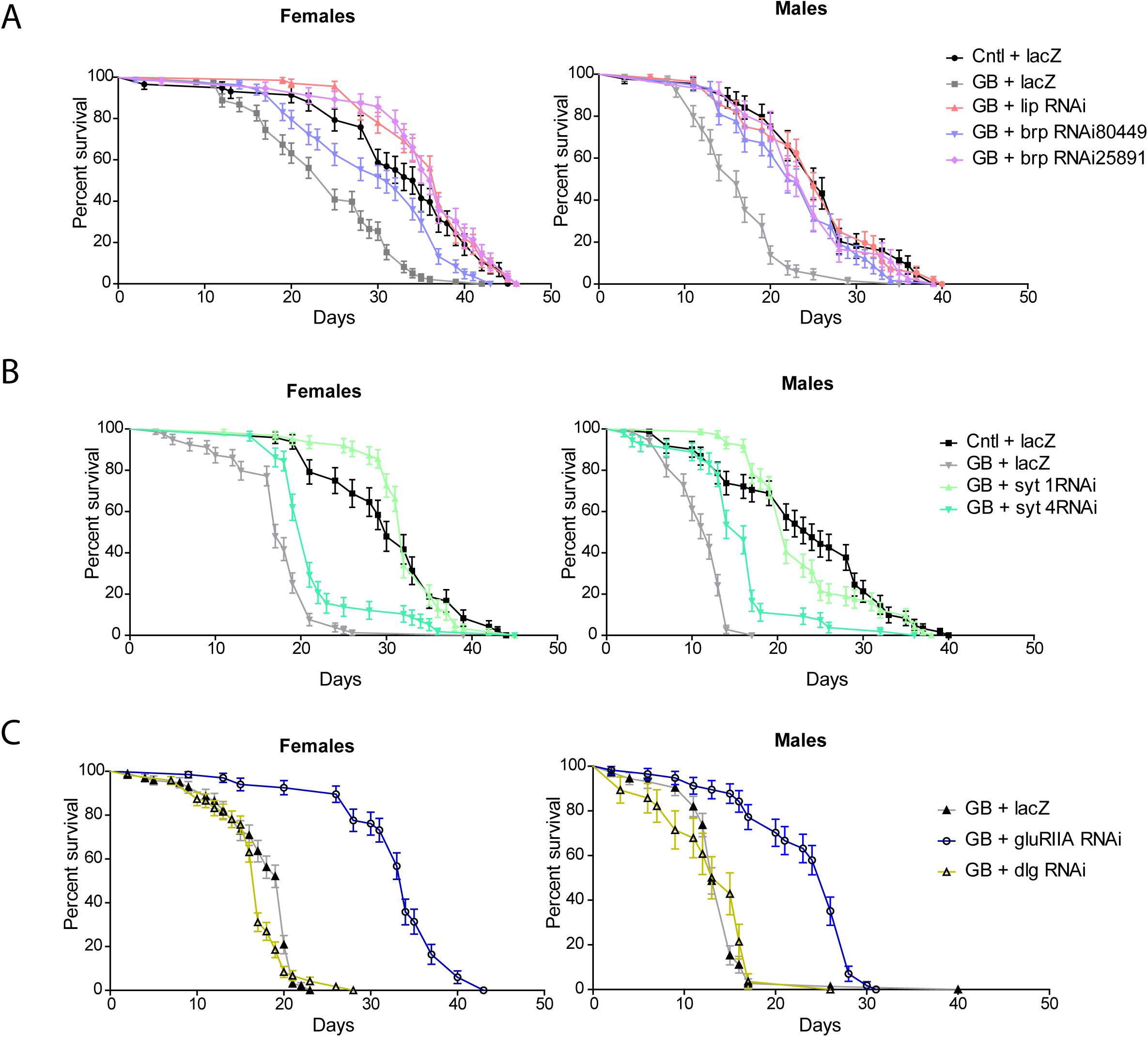
Synaptic genes contribute to premature death caused by glioblastoma. A) Graphs showing the percentage of survival of adult flies in control wt (black), GB control (grey), GB + *lip* α *RNAi* (pale orange) and GB + *Brp RNAi* with two different RNAi lines (blue and purple). Females (left) and males (right) were analyzed separately. B) Graphs show the percentage of survival of control wt flies (black), GB control (grey), GB + *Syt 1 RNAi* (bright green) and GB + *Syt 4 RNAi* (green). Females (left) and males (right) were analyzed separately. C) Graphs showing the percentage of survival of control GB control flies (grey), GB + *glyRIIA RNAi* (blue) and GB + *dlg RNAi* (yellow). Females (left) and males (right) were analyzed separately. Logrank Test (Mantel-Cox) for trend analysis.

**Figure 7.**
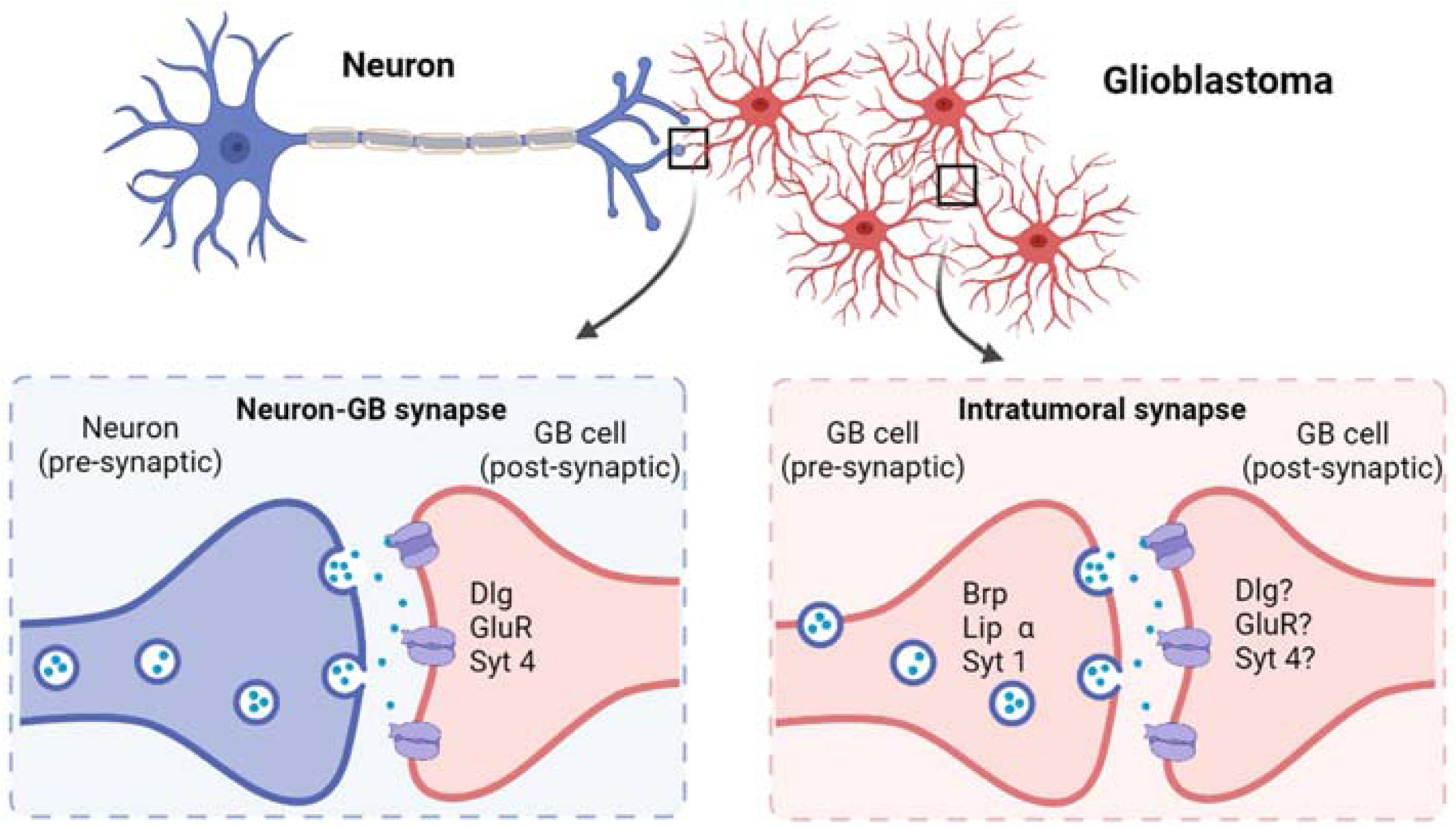
Summary. Schematic representation of neuron-GB synapse and intratumoral synapses. Bottom left, Neuron (blue) is the presynaptic component and GB cell (red) is the postsynaptic component. GB cells express postsynaptic genes *dlg, GluR* and *Syt 4*. Bottom right, intratumoral synapses are formed between two GB cells (red), presynaptic GB cells express *Brp, Lip* α and *Syt 1*. The other GB cells behave as postsynaptic, and have the same identity as in Neuron-GB synapses (*dlg, GluR* and *Syt 4)*. Created with BioRender.com

In addition, we analyzed the contribution of *GluRII, dlg* and *Syt 4* to life span in GB. The results show that *GluRIIA* or *Syt4* knockdown in GB cells, expands the life span of animals with GB, but the expression of *dlg* does not modify the premature death caused by GB progression (Figure 6 B-C).

## Discussion

In addressing the mechanisms that facilitate cell to cell communication in GB progression, we found genes that encode for synaptic proteins and are required for GB progression, suggesting a contribution of synapses in GB. Previous reports indicated the existence of glutamatergic synapses structured as neuron pre-synaptic, and GB cells post-synaptic. We have recapitulated neuron to GB synapses in *Drosophila melanogaster* and confirmed this unidirectional structure of glutamatergic synapses by GRASP experiments. We determined the expression and accumulation of post-synaptic proteins such as GluRII and Dlg in GB membranes, in the close area of Brp pre-synaptic protein. Moreover, the expression of post-synaptic genes including *GluRII, dlg and syt4* in GB cells is required for tumor progression, and for the deleterious consequences caused by GB, including neurodegeneration and premature death.

However, *dlg* knockdown in GB cells shows a particular phenotype, *dlg RNAi* prevents brain volume expansion and GB cells number increase, and also attenuates the synapse loss caused by GB. However it is not sufficient to prevent premature death caused in GB. It is tempting to discuss the contribution of Dlg to GB progression, and furthermore, the requirements of Dlg in glial cells for the normal function. Dlg protein is involved in post-synaptic structures, but also in cell polarity, neuronal differentiation and organization, and septate junctions in cellular growth control during larval development. In addition, Dlg contains a guanylate kinase domain that suggests a role in cell adhesion and signal transduction to control cell proliferation (Albertson and Doe, 2003; Koh et al., 1999; Li et al., 2009; Maiya et al., 2012; Ohshiro et al., 2000; Zhang et al., 2007). In consequence, *dlg* knockdown could affect a number of cellular functions that reduces life span, independently of the prevention of GB progression.

The multiple functions of many proteins has recently emerged as a novel point of view in biology, and reconciles the complex mechanisms behind cellular physiology, and the limited number of known genes. For example, Troponin-I was described as a central player in muscle formation, but recent discoveries show that Troponin-I is also involved in apico-basal polarity, chromosome stability and tumorigenesis (Casas-Tintó and Ferrús, 2019; Casas-Tintó et al., 2016; Sahota et al., 2009). Moreover, Caspases are another example of multi-functional proteins involved not only in apoptosis, but also in cancer progression and quiescence (Arthurton et al., 2020; Baena-Lopez, 2018; Baena-Lopez et al., 2018). Therefore, the impact of Dlg in GB and viability goes in line with this multiple functions of proteins, and brings a complex scenario that will require further study.

### Pre-synaptic genes are required for GB progression

In addition, we investigated the contribution of synaptic genes that encode for pre-synaptic proteins, such as Lip α, Syt 1 and Brp. Lip α and Syt 1 appeared as hits in the unbiased genetic screening that we performed to search for anti-GB strategies, suggesting a contribution for pre-synaptic genes in GB. However, GB does not form any pre-synaptic structure with regards to the synapses established with neurons according to GRASP results. Therefore, we discarded that the pre-synaptic components in GB are related to GB-neuron synapses. In consequence, we propose for the first time that GB cells establish intratumoral synapses and these are required for GB aggressiveness.

In particular, we have evaluated the contribution of Syt 1, Lip α and Brp pre-synaptic proteins. We found that all of them contribute to calcium signaling enhancement in GB. Additionally we demonstrated that GB cells upregulate Syt 1, determined by the accumulation of a Syt1-GFP fusion protein in GB tissue. The expression of a tagged Lip-GFP form in GB cells shows an increase in the dotted signal compatible with the formation of presynaptic structures in GB cells. These pieces of evidence support the formation of intratumoral synapses. We also obtained high resolution TEM images that show dense structures between GB cells, comparable to synaptic densities. These structures are morphologically disrupted upon *Syt1* or *lip* α knockdown in GB, supporting the hypothesis of GB-GB synapses. Finally, *brp* or *lip* α RNAi do not prevent GB cells number increase nor tumor volume, however they do prevent synapse loss in neurons and life span shortening indicating that intratumoral synapses contribute to neurodegeneration and lethality more than to GB expansion. Altogether these evidences support the presence and contribution of intratumoral synapses to GB progression.

In spite of all these results, we cannot determine if intratumoral synapses function as glutamatergic synapses described in the nervous system. Our results indicate that synaptic proteins are relevant for calcium waves, which are associated to activity in glial cells, GB progression and negative consequences, and we have observed the accumulation of proteins comparable to functional synapses, but the precise mechanisms that underlie intratumoral synapses will have to be determined in the future. It is proposed that synaptic proteins have ancestral functions related to secretion of neurotransmitters in chonoflagellates, and conserved through metazoans. Moreover, studies on postsynaptic proteins in choanoflagellates revealed unexpected localization patterns and new binding partners, both which are conserved in metazoans (Alié and Manuël, 2010; Burkhardt, 2015). New alternative functions of synaptic structures or synaptic proteins are possible and might not imply the transmission of impulses between two cells. There is evidence that indicates a coordination among glial cells under normal conditions, and in GB (Osswald et al., 2015), and we have observed a reduction in the calcium influx in GB cells upon presynaptic and postsynaptic genes knockdown.

Moreover, GB progression relies on the formation of cytoneme-like structures, tumor microtubes (Casas-Tintó and Portela, 2019; Osswald et al., 2015; Portela et al., 2019b). Cytonemes in epithelial cells accumulate receptors that contribute to cell-to-cell communication. In particular, components of neuronal synapses function in proper cytoneme formation and signaling including GluR, Syt 4 and Syt 1 (Huang et al., 2019). Thus it is possible that by downregulating synaptic proteins we were disrupting cytoneme formation and hence preventing GB growth.

On the other hand, breast to brain metastasis is driven by activation of N-methil-D-aspartate receptors (NMDAR) through glutamate ligands. Metastatic tumor cells do not produce sufficient glutamate ligands to induce signaling, which is achieved by the formation of tripartite synapses between cancer cells and neurons (Zeng et al., 2019). Besides it has been shown that samples of human cancers such as gastrointestinal stromal tumors (Bu□mming et al., 2007), have enriched synaptic proteins, thus it is plausible to hypothesize that synaptic proteins overexpression could be a common mechanism for cancer progression.

In conclusion, the results presented in this manuscript open new avenues to understand the mechanisms that coordinate the communication between GB cells, and the relation with the surrounding healthy cells and other tumoral cells. The strategies developed by neuroscientists during the last decades to modulate synapse number or activity emerge as a promising possibility to modulate the progression of GB, and maybe other tumors of the nervous system. The potential of these novel targets to prevent GB expansion deserve further investigation in line with the relevance of synaptic coordination within the tumoral mass.

